# The prion protein is not required for peripheral nerve repair after crush injury

**DOI:** 10.1101/2020.10.20.348011

**Authors:** Anna Henzi, Adriano Aguzzi

## Abstract

The cellular prion protein (PrP) is essential to the long-term maintenance of myelin sheaths in peripheral nerves. PrP activates the adhesion G-protein coupled receptor Adgrg6 on Schwann cells and initiates a pro-myelination cascade of molecular signals. Because Adgrg6 is crucial for peripheral myelin development and regeneration after nerve injury, we investigated the role of PrP in peripheral nerve repair. We performed experimental sciatic nerve crush injuries in co-isogenic wild-type and PrP-deficient mice, and examined peripheral nerve repair processes. Generation of repair Schwann cells, macrophage recruitment and remyelination were similar in PrP-deficient and wild-type mice. We conclude that PrP is dispensable for sciatic nerve regeneration after crush injury. Adgrg6 may sustain its function in peripheral nerve repair independently of its activation by PrP.

## Introduction

The reciprocal interaction between axons, Schwann cells and the extracellular matrix is crucial for development, maintenance and repair of the peripheral nervous system (PNS) (1,2). The adhesion G-protein coupled receptor Adgrg6 (formerly called Gpr126) was shown to be required for Schwann cell development (3), peripheral nerve repair (4,5) and possibly myelin maintenance (4). Three natural ligands for Adgrg6 have been described: collagen IV (6), laminin-211 (7) and the prion protein (PrP) (8). Collagen IV and laminins are essential components of the Schwann cell basal lamina, which plays an important role in developmental myelination and remyelination after injury (9–11). PrP is a glycophosphoinositol-anchored glycoprotein highly expressed in the nervous system (12) and mainly known for its role in prion diseases (13,14). The development of chronic peripheral demyelinating neuropathy in PrP knockout mice has led to the identification of cellular PrP as necessary for myelin maintenance (15,16). This finding raises the question whether PrP may also play a role in PNS repair. Here, we hypothesized that PrP may represent one of the ligands required to activate Adgrg6 after nerve injury. If so, PrP-deficient mice might develop similar defects in nerve regeneration as were reported in Adgrg6 knockout mice. To address this hypothesis, we performed nerve crush injuries in female *Prnp* knockout mice (*Prnp*^ZH3/ZH3^) and compared the morphological and molecular stages of repair to wild type (WT) mice. Contrary to our expectations, we detected no differences in peripheral nerve repair between *Prnp*^ZH3/ZH3^ and WT mice. These findings suggest that PrP is dispensable for peripheral nerve repair.

## Material and methods

### Mice

Breeding and maintenance of mice was performed in laboratory animal facilities (optimal hygienic conditions)at the University Hospital Zurich. Mice were housed in groups of 3 with unlimited access to food and water. Animal care and experiments were performed in accordance with the Swiss Animal Protection Law. All experiments were approved by the Veterinary Office of the Canton of Zurich (permit ZH168/2019).

### Nerve crush surgery

Surgery was performed on the right hind limb of two-month-old female WT and *Prnp*^ZH3/ZH3^ mice as described previously (17,18). Briefly, mice were injected with buprenorphinum (Temgesic, 0.1 mg/kg of bodyweight) prior to surgery. The surgery was performed under isoflurane anaesthesia. Fur was removed with an electric trimmer and the sciatic nerve was exposed at the height of the hip by making a small incision. The sciatic nerve was exposed by blunt dissection and crushed by squeezing firmly for 30 seconds with a Dumont S&T JF‐5 forceps (FST tools) at the height of the sciatic notch. The tissue was repositioned, and the wound was sealed with a suture. Mice were administered buprenorphinum for analgesia during the first 2 postoperative days. Sciatic nerves from crushed and contralateral side were harvested at 5, 10, 12, 16 and 30 days post crush.

### Nerve harvesting

Mice were sacrificed by cervical dislocation in deep anaesthesia. Sciatic nerves were embedded in the required fixation solution for morphological analysis or frozen in liquid nitrogen for protein analysis. After crush injury, the sciatic nerve distal to the crush site was divided in 3 parts (see Supplementary Fig. 1). A segment distal to the crush site (3 mm) was used for electron microscopy (EM). The distal side of this segment was embedded facing the front of the block face for sectioning. Sections from 2 mm distal to the crush site were analysed by EM. The remaining sciatic nerve was cut in half and the more proximal segment was processed for immunofluorescence (IF) as described below. the nerve segment was positioned with the proximal side facing the front of the block face for sectioning. Thereby, the EM and IF images derived from a similar distance from the crush side. The most distal segment of the sciatic nerve was frozen in liquid nitrogen and used for protein analysis. On the contralateral side, the corresponding segments were collected in the same manner.

### Morphological analysis by EM

Sciatic nerves were immersed in 4% glutaraldehyde in 0.1 M sodium phosphate buffer pH 7.4 immediately after dissection and incubated at 4 °C overnight. Tissue was embedded in Epon using standard procedures. Further steps were performed as described (18). Briefly, 99 nm sections were collected on ITO coverslips (Optics Balzers). Imaging for EM reconstruction of the entire sciatic nerve section was performed with either a Carl Zeiss Gemini Leo 1530 FEG or Carl Zeiss Merlin FEG scanning electron microscope attached to Atlas modules (Carl Zeiss). Adobe Photoshop CS5 was used for image analysis. The g-ratio corresponds to the ratio between axon diameter and fibre diameter. The axon diameter was derived from the axon area. The myelin thickness was measured at two different locations of the myelin ring. The average myelin thickness was added twice to the axon diameter to obtain the fibre diameter. For g-ratio quantification, three different locations on the cross section were chosen and at least 100 fibres per sample were analysed. The number of intact appearing myelin profiles, remyelinated fibres and the area covered by myelin debris was assessed manually on the entire cross section. The investigator was blinded as to the genotype of the mice for all analyses.

### Immunofluorescence (IF)

Sciatic nerves were isolated as described above. The tissue was fixed in 4% paraformaldehyde overnight at 4°C. Then, the tissue was incubated in 30% sucrose solution and frozen in OCT compound. Cross-sections of 8 µM thickness were cut at the cryostat. After drying at room temperature (RT), sections were incubated in blocking buffer (10% normal goat serum, 0.5% bovine serum albumin (BSA), 0.3% Triton X-100 in PBS) for 1 h at RT. Blocking buffer was diluted 1:1 with PBS for incubation with primary antibodies at 4 °C overnight. The following primary antibodies were used: Ki-67 (Abcam, ab15580, 1:1000), rabbit anti-c-Jun (Cell Signalling Technologies, 9165, 1:250), mouse anti-S100 (Dako, Z0311, 1:250), rat anti-CD68 (Biorad, MCA1957, 1:200). After three washes in PBS, slides were incubated with fluorophore-conjugated secondary antibodies (Alexa Fluor goat-anti rabbit 488 nm, goat anti-rat 594 nm, Invitrogen) for 1 h at RT. Then, sections were washed in PBS, incubated with DAPI (1:10’000) to label nuclei for 10 min and mounted using Fluorescence mounting medium (Dako, S3023). Fluorescent images were recorded with a Leica SP5 Confocal Microscope.

For quantification of IF images, ImageJ (version 1.52) was used. To count DAPI, c-Jun and Ki67-positive nuclei, nuclei were defined using the threshold function. Counting was performed on binary images. For each marker, the same ImageJ macro was applied to all images from one time point. The number of Ki67-positive or c-Jun-positive nuclei was normalized to the total number of DAPI-positive nuclei in the field of view. Macrophages were quantified as the total CD68-positive area in relation to the S100-positive area. For each time point, three animals per group were analysed. For each mouse, the mean value from quantification of 2-4 images from separate nerve sections was used for statistical analysis. The investigator was blinded as to the time point and the genotype of the mice.

### Western blot analysis

Sciatic nerves were homogenized using stainless-steel beads in ice-cold lysis buffer (phosSTOP (Sigma, 4906845001) and protease inhibitor (Sigma, 11836153001) in RIPA buffer). Debris was removed by centrifugation of the lysates for 10 min at 10’000 g. Protein concentration was measured with BCA assay (Thermo Scientific). 10 μg protein for each sample was boiled in 4 × LDS (Invitrogen) at 95 °C for 5-10 min and loaded on a gradient 4-12% NuPAGE Bis-Tris gels (Invitrogen). After electrophoresis at 200 V, gels were transferred to PVDF membranes with the iBlot system (Life Technologies). Then, membranes were blocked with 5% SureBlock (LuBioScience GmbH, SB232010) in TBS-T. Incubation with primary antibodies was performed night at 4 °C. After three washes for 10 min, membranes were incubated with secondary antibodies coupled to horseradish peroxidase for 1 h at room temperature (RT). Membranes were then washed and developed with Crescendo chemiluminescence substrate system (Sigma, WBLUR0500). Signals were detected using a Fusion Solo S imaging system (Vilber). The FusionCapt Advance software was used for densitometry. Each lane in the blots and each point in the graphs represents one sciatic nerve from one mouse. Original, uncropped images are shown in supplementary figure 2. The following primary antibodies were used for western blotting: GFAP (1:2000, Cell Signaling Technologies, 12389S), c-Jun (1:1000, 9165s), Calnexin (1:2000, Enzo, ADI-SPA-865-D). In addition, we used an in-house produced mouse monoclonal antibody (POM2) for detection of PrP (19). The following horseradish peroxidase coupled secondary antibodies were used: anti-mouse IgG (1:10’000, Jackson Immuno Research, 115-035-003), anti-rabbit IgG (1:4000, Jackson Immuno Research, 111-035-003).

### Experimental design and statistical analysis

Statistical analysis was performed with GraphPad Prism software (version 8.4.2). We assumed normal distribution and equal variances of data but did not formally test this assumption due to small *n* values. Unpaired two-tailed t-test was used for comparison of two groups. For comparison of three or more groups, two-way ANOVA followed by Sidak’s multiple comparison test was used and multiplicity adjusted p-values were reported. P-values below 0.05 were considered statistically significant. P-values are indicated in graphs as *: *p* < 0.05. ns: not significant, *p* > 0.05. Error bars in graphs show SEM. No samples or data were omitted during the analyses. R (version 3.5.2) was used to generate the scatterplot visualizing the g-ratio analysis.

## Results

### PrP is not required for demyelination, repair Schwann cell generation and proliferation following nerve injury

To study the role of PrP in peripheral nerve regeneration, we performed sciatic nerve crush injuries in female *Prnp*^ZH3/ZH3^ and WT mice at 2 months of age and investigated the morphological and molecular stages of peripheral nerve repair. We crushed the sciatic nerve at the sciatic notch, and harvested the nerve segments distal to the injury site at 5, 10, 12, 16 and 30 days post crush (d.p.c). Additionally, we collected the uninjured contralateral sciatic nerves as controls. To ensure comparability between the groups, the sections harvested for the various analyses were taken from the same distance from the crush site as depicted in supplementary figure 1.

First, we assessed if Schwann cells in *Prnp*^ZH3/ZH3^ mice were able to properly transdifferentiate to repair Schwann cells. We investigated c-Jun as a molecular marker of repair Schwann cells (20) by immunofluorescence (IF) and western blotting (Fig. 1a-d). At 5 d.p.c. a strong upregulation of c-Jun was detected both in WT and *Prnp*^ZH3/ZH3^ mice. c-Jun levels decreased at later time points post injury. In uninjured contralateral nerves, c-Jun protein levels were low and nuclear c-Jun staining in IF was very faint. We did not detect any significant differences in c-Jun levels between WT and *Prnp*^ZH3/ZH3^ mice at 5, 10, 16 and 30 d.p.c.

**Figure 1:**
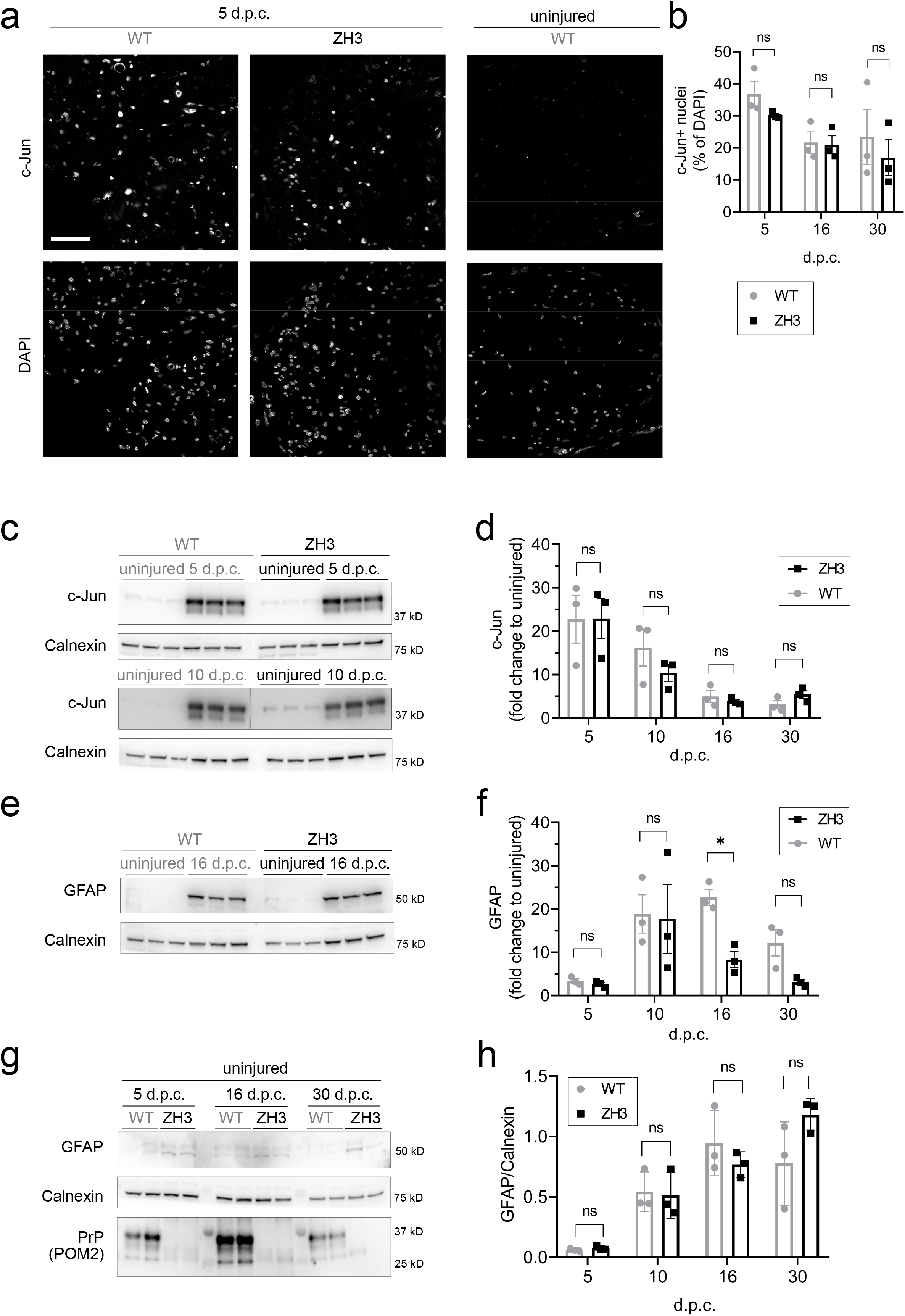
Generation of repair Schwann cells. **(a)** IF images of sciatic nerve cross sections from WT and *Prnp*^ZH3/ZH3^ mice at 5 and 10 d.p.c. showed upregulation of c-Jun. Nuclear c-Jun expression was faint in the uninjured sciatic nerve of a WT mouse. Channels for c-Jun and DAPI are shown separately. Scale bar 50 μM. **(b)** C-Jun upregulation as quantified from IF images was highest at 5 d.p.c. Quantification of c-Jun positive nuclei did not reveal significant differences between WT and *Prnp*^ZH3/ZH3^ mice at 5, 16 and 30 d.p.c (two-way ANOVA, Sidak’s multiple comparisons test, *p* > 0.05). **(c)** Western blot of lysates from crushed and uninjured (contralateral) sciatic nerves at 5 and 10 d.p.c. showed a pronounced upregulation of total c-Jun protein levels both in WT and *Prnp*^ZH3/ZH3^ mice. **d)** Quantification of c-Jun upregulation from western blots revealed no significant difference between WT and *Prnp*^ZH3/ZH3^ mice at the investigated time points (two-way ANOVA, Sidak’s multiple comparisons test, *p* > 0.05). For each mouse (*n* = 3 per genotype and time point), the ratio of c-Jun in the crushed sciatic nerve compared to the uninjured side was calculated. **(e)** Western blot of lysates from crushed and uninjured (contralateral) sciatic nerves at 16 d.p.c. showed a strong upregulation of GFAP both in WT and *Prnp*^ZH3/ZH3^ mice. **(f)** Quantification of GFAP by densitometry (injured nerve vs. contralateral side) showed a slightly stronger upregulation in WT mice than in in *Prnp*^ZH3/ZH3^ mice, which reached statistical significance at 16 d.p.c. as assessed by two-way ANOVA followed by Sidak’s multiple comparisons test (*p* = 0.00425 at 16 d.p.c., *p* > 0.05 at 5, 10 and 30 d.p.c.). **(g)** Western blot detecting low GFAP levels in uninjured sciatic nerves from WT (*n* = 6) and *Prnp*^ZH3/ZH3^ mice (*n* = 6) sacrificed at 5, 16 and 30 d.p.c. GFAP levels were higher in *Prnp*^ZH3/ZH3^ sciatic nerves when compared to WT nerves. Blotting with POM2, a monoclonal anti-PrP antibody, detected PrP in WT but not in *Prnp*^ZH3/ZH3^ sciatic nerves, confirming the absence of PrP in *Prnp*^ZH3/ZH3^ mice. **(h)** The absolute GFAP levels in sciatic nerves after crush were not significantly different when comparing *Prnp*^ZH3/ZH3^ and WT mice (*p* > 0.05 at all time points as assessed by two-way ANOVA followed by Sidak’s multiple comparisons test). n.s. = not significant.

GFAP, another marker of repair Schwann cells, peaked one week later than c-Jun at 10-16 d.p.c (Fig. 1e,f). Quantification of GFAP upregulation after crush (Fig. 1f) revealed lower GFAP upregulation in *Prnp*^ZH3/ZH3^ mice compared to WT mice, which reached the threshold for statistical significance at 16 d.p.c. (*p* = 0.0425). This differential upregulation might be related to the fact that *Prnp*^ZH3/ZH3^ mice had on average a two-fold higher baseline GFAP level when compared to WT mice (Fig. 1g, relative increase to WT GFAP level as quantified from western blot: WT 1.00 ± 0.17 (*6*); *Prnp*^ZH3/ZH3^ 2.05 ± 0.09 (*6*); mean ± SEM (*n*); *p* = 0.0003, unpaired t-test). Compatible with this explanation, the absolute GFAP protein levels after nerve crush as assessed by western blotting were similar in WT and *Prnp*^ZH3/ZH3^ mice at all investigated time points (Fig. 1h). In conclusion, *Prnp*^ZH3/ZH3^ mice required less GFAP upregulation to reach WT levels after nerve crush due to their higher baseline levels.

Ki67 staining in crushed sciatic nerves showed that proliferation was highest at 5 d.p.c. both in WT and *Prnp*^ZH3/ZH3^ mice, and no significant difference in the proliferation index was detected between WT and *Prnp*^ZH3/ZH3^ mice at any of the time points investigated (Fig. 2a,b).

**Figure 2:**
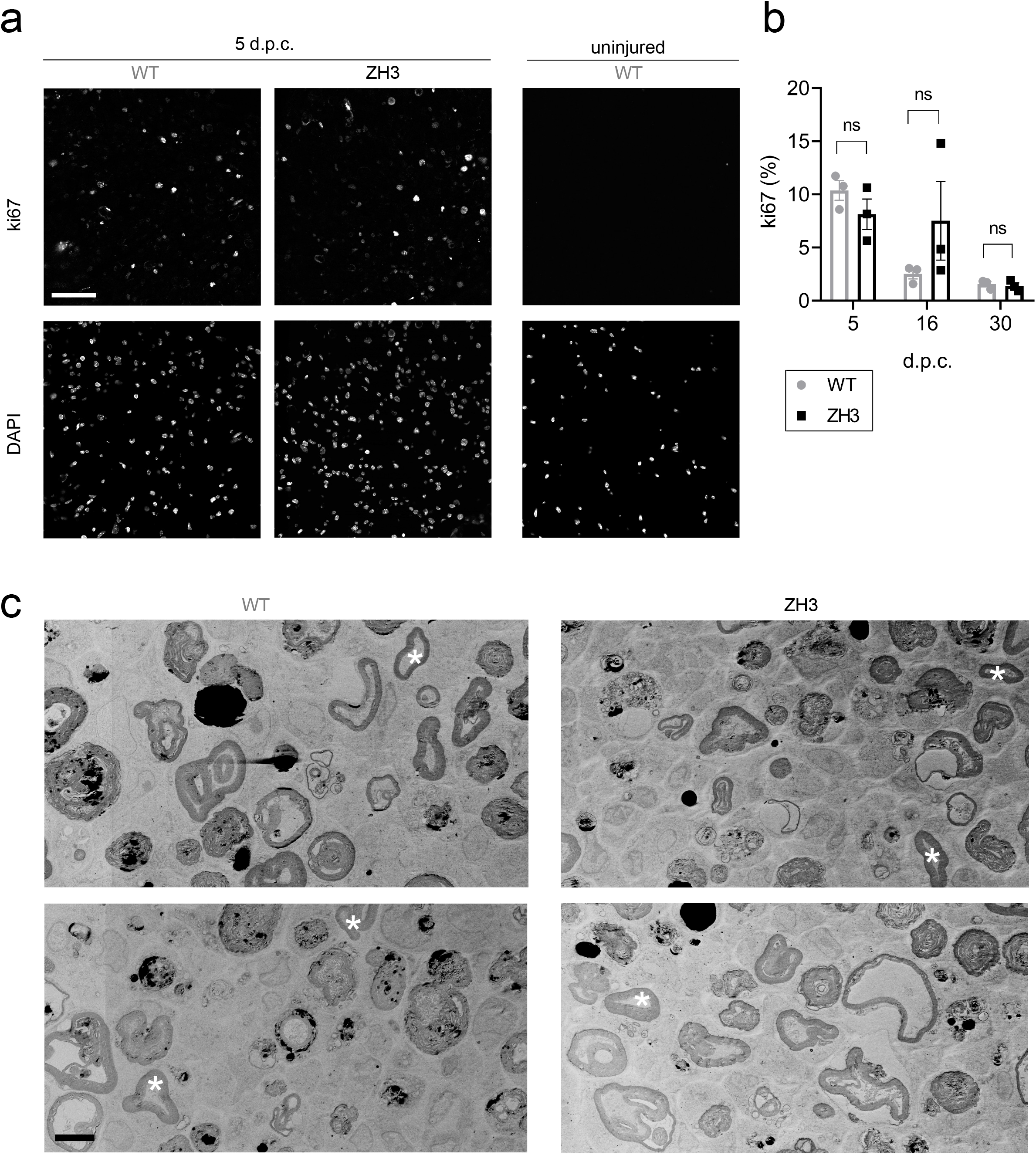
Proliferation and demyelination in the early stages after crush injury. **(a)** IF images of sciatic nerve cross sections from WT and *Prnp*^ZH3/ZH3^ mice at 5 d.p.c. revealed many ki67-positive proliferating cells. The contralateral uninjured nerve from a WT mouse showed no proliferating cells. Channels for ki67 and DAPI are shown separately. Scale bar: 50 μM. **(b)** Ki67-proliferation index (% ki67-positive nuclei) as quantified from IF images was highest at 5 d.p.c. and decreased at later stages post crush injury. No significant differences were detected between WT and *Prnp*^ZH3/ZH3^ mice at 5, 16 and 30 d.p.c. (two-way ANOVA, Sidak’s multiple comparisons test, *p* > 0.05). n.s. = not significant. **(c)** EM images of sciatic nerves from WT and *Prnp*^ZH3/ZH3^ mice at 5 d.p.c. showed extensive demyelination and few intact appearing myelin rings (white asterix). Scale bar: 5 μm.

In the early stages after nerve crush, Schwann cells are the main effectors of myelin breakdown and digestion (21). Using electron microscopy (EM) we investigated the extent of demyelination by counting the number of myelin profiles that still appeared intact at 5 d.p.c. (Fig. 2c). The number of intact appearing myelin profiles per sciatic nerve cross section was not significantly altered in *Prnp*^ZH3/ZH3^ mice when compared to WT mice (WT 191 ± 38 (*3*); *Prnp*^ZH3/ZH3^ 244 ± 68 (*3*); mean ± SEM (*n*); *p* = 0.5277; unpaired t-test), indicating that the initial myelin breakdown is neither slowed nor accelerated in the absence of PrP.

Collectively, these results suggested that PrP is not required for upregulation of repair Schwann cell markers, proliferation and early myelin breakdown after nerve crush injury.

### PrP is not required for macrophage recruitment after peripheral nerve injury

Blood derived macrophages are recruited to the injured nerve via chemokines secreted by Schwann cells and mesenchymal cells (22). Together with resident endoneurial macrophages, they contribute to debris clearance and repair after injury (23). We performed IF for the macrophage marker CD68 to investigate if PrP is playing a role in macrophage recruitment (Fig. 3a-b). While uninjured nerves were basically devoid of CD68-positive cells (Fig. 3b), crushed sciatic nerves were infiltrated by numerous macrophages (Fig. 3a). In both genotypes, macrophages were rounded and displayed a phagocytic, foamy appearance. We quantified the area covered by CD68-positive cells in relation to the S100-positive area (Fig. 3c). Macrophage accumulation in the endoneurium peaked at 10 d.p.c. in both genotypes, which is consistent with the time course of macrophage infiltration described in the literature (23,24). No differences in macrophage levels were detected between WT and *Prnp*^ZH3/ZH3^ sciatic nerves in this analysis, suggesting that the lack of PrP does not affect injury-induced macrophage recruitment to the peripheral nerves.

**Figure 3:**
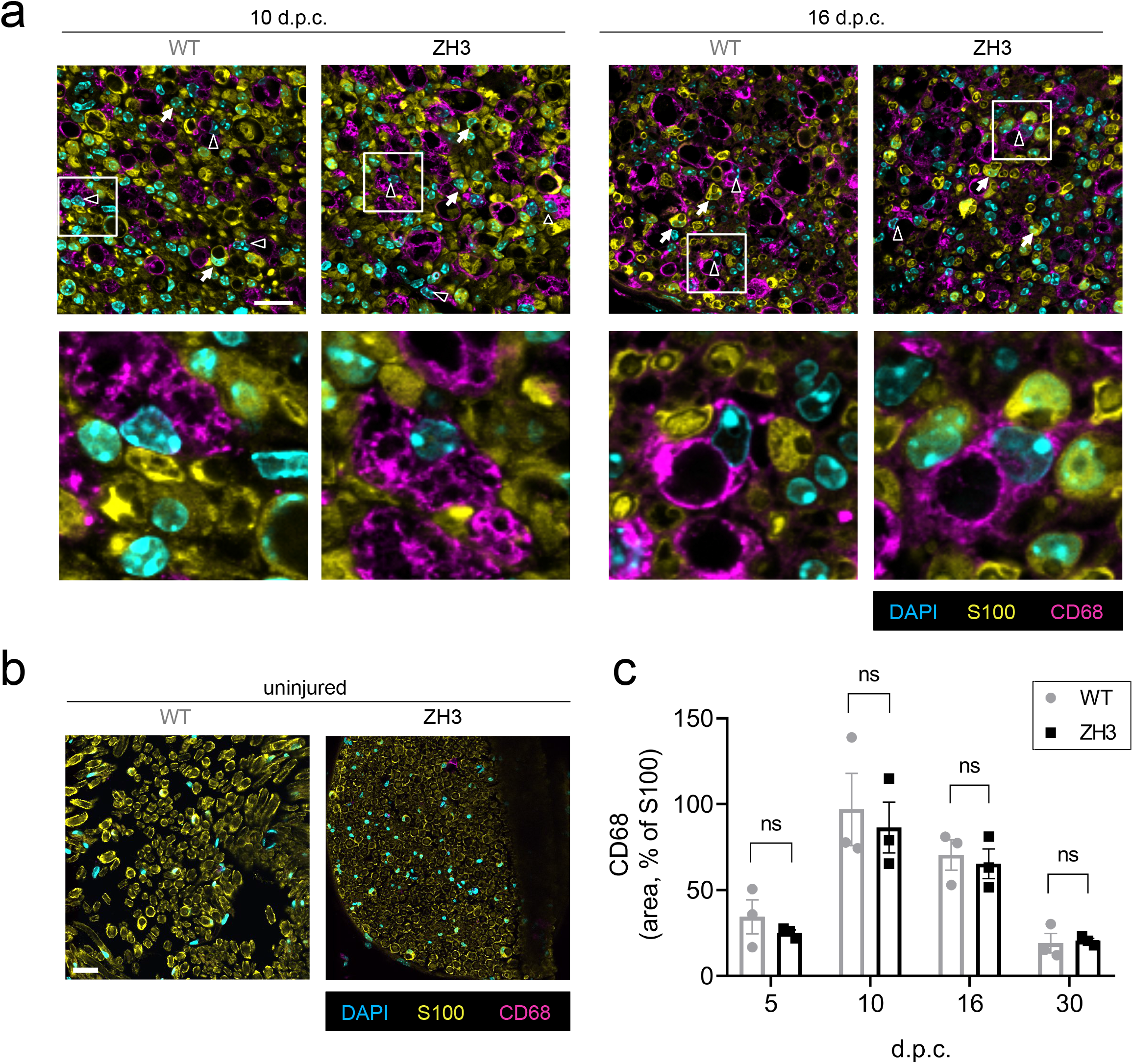
Recruitment of macrophages. **(a)** IF images of sciatic nerve cross sections from WT and *Prnp*^ZH3/ZH3^ mice at 10 and 16 d.p.c. showed pronounced infiltration by CD68-positive macrophages. White triangles: CD68-positive macrophages. White arrows: S100-positive Schwann cells. Squares indicate region selected for blown-up images of macrophages shown in the lower panels. Scale bar 25 μM. **(b)** IF images of uninjured (contralateral) sciatic nerve showed no infiltration by macrophages. Scale bar 25 μM. **(c)** Macrophage infiltration peaked at 10 d.p.c. both in WT and *Prnp*^ZH3/ZH3^ mice. The extent of macrophage infiltration (quantified as the ratio of the CD68-positive area to the S100-positive area in IF images) was not significantly different when comparing WT and *Prnp*^ZH3/ZH3^ mice at all investigated time points (two-way ANOVA, Sidak’s multiple comparisons test, *p* > 0.05). n.s. = not significant.

### Remyelination is not impaired in Prnp^ZH3/ZH3^ mice

To investigate if PrP is required in later stages of nerve repair, we assessed remyelination after nerve crush injury by EM. We first analysed the crushed sciatic nerves at 12 d.p.c., when regrown axons are undergoing remyelination by redifferentiated Schwann cells (Fig. 4a). EM analysis revealed robust remyelination in both WT and *Prnp*^ZH3/ZH3^ nerves with no statistically significant difference between genotypes (WT 85.88 ± 1.39 (*3*), *Prnp*^ZH3/ZH3^ 88.56 ± 1.28 (*3*), percent of remyelinated axons; mean ± SEM (*n*), *p* = 0.2282; unpaired t-test). No gross morphological differences were noted between genotypes. At 30 d.p.c. some myelin debris was still present both in WT and *Prnp*^ZH3/ZH3^ mice (Fig. 4b,e). The amount of remaining debris was similar in both genotypes (WT 2.62 ± 0.37 (*3*), *Prnp*^ZH3/ZH3^ 2.55 ± 0.29 (*3*), percentage of total sciatic nerve area covered by myelin debris; mean ± SEM (*n*), *p* = 0.8911; unpaired t-test). To determine the thickness of the myelin sheath, we performed g-ratio analysis at 10 and 30 d.p.c. In both genotypes, the g-ratio was higher in sciatic nerves at 10 d.p.c (WT 0.88 ± 0.002 (*335*), *Prnp*^ZH3/ZH3^ 0.88 ± 0.002 (*323*), g-ratio; mean ± SEM (*number of myelinated axons quantified*), *p* = 0.2745; unpaired t-test) than at 30 d.p.c. (WT 0.73 ± 0.003 (*319*), *Prnp*^ZH3/ZH3^ 0.72 ± 0.003 (*423*), g-ratio; mean ± SEM (*n*), *p* = 0.4419; unpaired t-test), indicating an increase in myelin thickness. There was no significant difference in myelin thickness as assessed by g-ratio analysis between WT and *Prnp*^ZH3/ZH3^ mice at either of the two time points (quantification for 30 d.p.c. shown in Fig. 4c,d).

**Figure 4:**
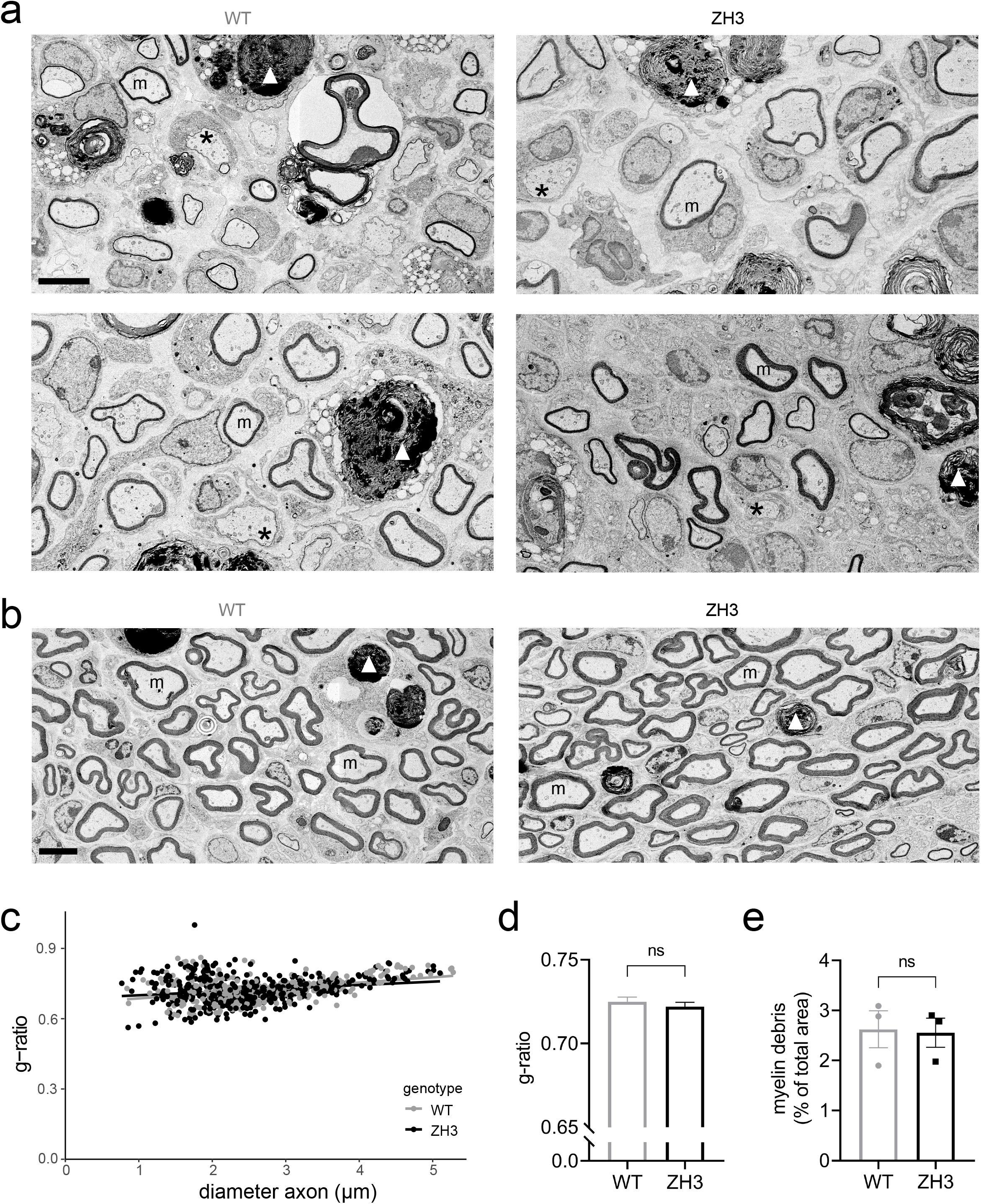
Remyelination. **(a)** EM images of sciatic nerves from WT and *Prnp*^ZH3/ZH3^ mice at 12 d.p.c. revealed robust remyelination in both genotypes. Scale bar 5 μm. **(b)** EM images of sciatic nerves from WT and *Prnp*^ZH3/ZH3^ mice at 30 d.p.c. showed remyelinated axons and little myelin debris remaining. m: myelinated axons, *: non-myelinated axon with > 1 μm diameter, white arrowhead: myelin debris. Scale bar 5 μm. **(c, d)**G-ratio analysis in WT (*n* = 3) and *Prnp*^ZH3/ZH3^ (*n* = 3) mice at 30 d.p.c. Quantification of > 100 myelinated axons per mouse. Straight lines in **(c)**show linear regression. No significant difference in mean g-ratio **(d)** was detected between genotypes (unpaired t-test, *p* > 0.05). **e)** Quantification of area covered by myelin debris at 30 d.p.c. showed no significant difference between WT (*n* = 3) and *Prnp*^ZH3/ZH3^ (*n* = 3) mice (unpaired t-test, *p* > 0.05). n.s. = not significant.

## Discussion

Adgrg6 is expressed in Schwann cells and was shown to have autonomous and non-autonomous functions in peripheral nerve repair (4,5). PrP is an agonist of Adgrg6 on Schwann cells (8). We therefore hypothesized that the lack of PrP causes similar defects in peripheral nerve repair as the knockout of Adgrg6. To evaluate our hypothesis, we performed sciatic nerve crush injuries in female WT and *Prnp*^ZH3/ZH3^ mice and investigated the stages of repair by EM, IF and western blotting. However, we found that peripheral nerve repair was not altered in *Prnp*^ZH3/ZH3^ mice when compared to WT mice. The extent and time point of c-Jun upregulation, demyelination, Schwann cell proliferation, macrophage recruitment and remyelination were similar in *Prnp*^ZH3/ZH3^ mice when compared to WT mice at all investigated time points. Our results thus suggest that PrP is dispensable for peripheral nerve repair. This may be either because PrP plays no major role in the peripheral nerve repair processes, or because other molecules can compensate for the lack of PrP. Alternatively, Adgrg6 may exhibit a basal constitutive activity sufficient to guarantee nerve repair in the absence of any agonist. Regarding the first point, PrP is dispensable for normal development of the PNS and only required for long-term maintenance of the myelin sheath (15). Many of the repair processes after nerve injury recapitulate developmental processes (25) and it is possible that PrP is simply not required for peripheral nerve repair. As to the second point, PrP is thought to function in myelin maintenance via activation of Adgrg6 on Schwann cells. Adgrg6 is involved in peripheral nerve repair, as was suggested by deficits in macrophage recruitment and remyelination after peripheral nerve injury in conditional Adgrg6 knockout mice. Thus, Adgrg6 must sustain its function in nerve repair independent of PrP, for example via activation by other ligands such as collagen IV (6) and laminin-211 (7).

In a further possible scenario, PrP might play contrasting roles in different cells types during the repair processes. Theoretically, the effects of PrP in different cell types may cancel each other out in a transgenic mouse with global PrP knockout as was used in the present study. However, considering that the peripheral nerve repair processes are regulated by a highly orchestrated, balanced interaction between many cell types, we feel that this is an unlikely scenario.

We have previously described that *Prnp*^ZH3/ZH3^ mice exhibit an early upregulation of repair Schwann cell markers in sciatic nerves, which probably represents an early sign of the peripheral nerve disease (26). In the present study, we confirmed that the 2-3 months old *Prnp*^ZH3/ZH3^ mice undergoing crush injury showed a two-fold upregulation of GFAP in their uninjured sciatic nerves when compared to WT mice. The absolute GFAP levels after crush injury were similar in both genotypes, and the elevated baseline GFAP level in *Prnp*^ZH3/ZH3^ mice did not interfere with regeneration. Whereas defective Schwann cell proliferation and delayed regeneration after nerve crush injury have been described in GFAP knockout mice (27,28), the effect of GFAP overexpression on PNS development, maintenance or repair has not been investigated (29,30). GFAP gain-of-function mutations cause Alexander disease, a rare astrogliopathy characterized by accumulation of GFAP in the central nervous system (31). Development of a peripheral demyelinating neuropathy was reported in an 8-year-old patient with Alexander disease (32), but apart from this case, there is currently no compelling evidence that GFAP accumulation or overexpression contributes to or causes PNS degeneration. The finding that sciatic nerve repair is not impaired in *Prnp*^ZH3/ZH3^ mice despite increased baseline GFAP levels might be interpreted as an indication that GFAP overexpression is compatible with peripheral nerve regeneration.

In conclusion, the molecular and morphological analyses presented here consistently show that PrP is dispensable for peripheral nerve repair. While this finding disproves the hypothesis that PrP is required to activate Adgrg6-mediated macrophage recruitment and remyelination after nerve crush injury, it does not rule out an ancillary, non-essential role for such interactions. Furthermore, it is possible that additional, hitherto undiscovered Adgrg6 ligands exist. The importance of such ligands to peripheral-nerve repair may vary in different species. Therefore, the negative results reported here should not discourage researchers from exploring Adgrg6 activation as a possible adjuvant therapy for peripheral nerve repair.

## Acknowledgements

AA is the recipient of a Swiss National Research Foundation Sinergia grant (CRSII5 183563), of an individual SNRF grant, and of a Distinguished Scientist Award of the Nomis Foundation. We thank Dr. Jorge A. Pereira for teaching, supervision and technical help with EM imaging, and Alexander Henzi for support with R programming and statistical analyses.

## Author’s contributions

AH: Conceptualization, Methodology, Investigation, Data curation and analysis, Writing – original draft, review and editing. AA: Conceptualization, Data analysis, Resources, Supervision, Funding acquisition, Project administration, Writing – review and editing.

## Supplementary Figures

**Supplementary Figure S1: Nerve harvest after crush injury.** The sciatic nerve was crushed using a forceps at the sciatic notch. For harvesting, the nerve was cut at 3 mm distal to the crush side. The proximal segment was embedded for electron microscopy (EM), the distal segment was used for immunofluorescence (IF) and western blotting.

**Supplementary Figure S2: Original western blot and immunofluorescence images.** The uncropped blots have been inverted using the Affinity Photo software. The specific band is marked with * when additional non-specific bands are present. No lanes were cropped for presentation of the blots in the figures. The original immunofluorescence images are shown in black and white for separate channels (ki67 and c-Jun) or pseudo colours (merged, for CD68) with no alterations of contrast or brightness.

## References

1. Monk KR, Feltri ML, Taveggia C. New insights on schwann cell development: Schwann Cell Development. Glia. 2015 Aug;63(8):1376–93.

2. Raphael AR, Talbot WS. New Insights into Signaling During Myelination in Zebrafish. In: Current Topics in Developmental Biology [Internet]. Elsevier; 2011 [cited 2020 Jun 26]. p. 1–19. Available from: https://linkinghub.elsevier.com/retrieve/pii/B9780123859754000073

3. Monk KR, Oshima K, Jors S, Heller S, Talbot WS. Gpr126 is essential for peripheral nerve development and myelination in mammals. Development. 2011 Jul 1;138(13):2673–80.

4. Jablonka-Shariff A, Lu C-Y, Campbell K, Monk KR, Snyder-Warwick AK. Gpr126/Adgrg6 contributes to the terminal Schwann cell response at the neuromuscular junction following peripheral nerve injury. Glia. 2019 Dec 24;

5. Mogha A, Harty BL, Carlin D, Joseph J, Sanchez NE, Suter U, et al. Gpr126/Adgrg6 Has Schwann Cell Autonomous and Nonautonomous Functions in Peripheral Nerve Injury and Repair. J Neurosci. 2016 07;36(49):12351–67.

6. Paavola KJ, Sidik H, Zuchero JB, Eckart M, Talbot WS. Type IV collagen is an activating ligand for the adhesion G protein-coupled receptor GPR126. Science Signaling. 2014 Aug 12;7(338):ra76–ra76.

7. Petersen SC, Luo R, Liebscher I, Giera S, Jeong S-J, Mogha A, et al. The Adhesion GPCR GPR126 Has Distinct, Domain-Dependent Functions in Schwann Cell Development Mediated by Interaction with Laminin-211. Neuron. 2015 Feb;85(4):755–69.

8. Küffer A, Lakkaraju AKK, Mogha A, Petersen SC, Airich K, Doucerain C, et al. The prion protein is an agonistic ligand of the G protein-coupled receptor Adgrg6. Nature. 2016 25;536(7617):464–8.

9. Chen Z-L, Strickland S. Laminin γ1 is critical for Schwann cell differentiation, axon myelination, and regeneration in the peripheral nerve. Journal of Cell Biology. 2003 Nov 24;163(4):889–99.

10. Wallquist W, Patarroyo M, Thams S, Carlstedt T, Stark B, Cullheim S, et al. Laminin chains in rat and human peripheral nerve: Distribution and regulation during development and after axonal injury. J Comp Neurol. 2002 Dec 16;454(3):284–93.

11. Yu W-M, Yu H, Chen Z-L, Strickland S. Disruption of laminin in the peripheral nervous system impedes nonmyelinating Schwann cell development and impairs nociceptive sensory function. Glia. 2009 Jun;57(8):850–9.

12. Bendheim PE, Brown HR, Rudelli RD, Scala LJ, Goller NL, Wen GY, et al. Nearly ubiquitous tissue distribution of the scrapie agent precursor protein. Neurology. 1992 Jan 1;42(1):149–149.

13. Prusiner S. Novel proteinaceous infectious particles cause scrapie. Science. 1982 Apr 9;216(4542):136–44.

14. Aguzzi A, De Cecco E. Shifts and drifts in prion science. Science. 2020 02;370(6512):32–4.

15. Bremer J, Baumann F, Tiberi C, Wessig C, Fischer H, Schwarz P, et al. Axonal prion protein is required for peripheral myelin maintenance. Nat Neurosci. 2010 Mar;13(3):310–8.

16. Sakaguchi S, Katamine S, Nishida N, Moriuchi R, Shigematsu K, Sugimoto T, et al. Loss of cerebellar Purkinje cells in aged mice homozygous for a disrupted PrP gene. Nature. 1996 Apr 11;380(6574):528–31.

17. Gökbuget D, Pereira JA, Opitz L, Christe D, Kessler T, Marchais A, et al. The miRNA biogenesis pathway prevents inappropriate expression of injury response genes in developing and adult Schwann cells. Glia. 2018 Dec;66(12):2632–44.

18. Pereira JA, Gerber J, Ghidinelli M, Gerber D, Tortola L, Ommer A, et al. Mice carrying an analogous heterozygous dynamin 2 K562E mutation that causes neuropathy in humans develop predominant characteristics of a primary myopathy. Human Molecular Genetics. 2020 May 28;29(8):1253–73.

19. Polymenidou M, Moos R, Scott M, Sigurdson C, Shi Y, Yajima B, et al. The POM Monoclonals: A Comprehensive Set of Antibodies to Non-Overlapping Prion Protein Epitopes. Joly E, editor. PLoS ONE. 2008 Dec 8;3(12):e3872.

20. Arthur-Farraj PJ, Latouche M, Wilton DK, Quintes S, Chabrol E, Banerjee A, et al. c-Jun Reprograms Schwann Cells of Injured Nerves to Generate a Repair Cell Essential for Regeneration. Neuron. 2012 Aug;75(4):633–47.

21. Gomez-Sanchez JA, Carty L, Iruarrizaga-Lejarreta M, Palomo-Irigoyen M, Varela-Rey M, Griffith M, et al. Schwann cell autophagy, myelinophagy, initiates myelin clearance from injured nerves. Journal of Cell Biology. 2015 Jul 6;210(1):153–68.

22. Toma JS, Karamboulas K, Carr MJ, Kolaj A, Yuzwa SA, Mahmud N, et al. Peripheral Nerve Single-Cell Analysis Identifies Mesenchymal Ligands that Promote Axonal Growth. eNeuro. 2020 May;7(3):ENEURO.0066-20.2020.

23. Mueller M, Leonhard C, Wacker K, Ringelstein EB, Okabe M, Hickey WF, et al. Macrophage Response to Peripheral Nerve Injury: The Quantitative Contribution of Resident and Hematogenous Macrophages. Lab Invest. 2003 Feb;83(2):175–85.

24. Taskinen HS, Röyttä M. The dynamics of macrophage recruitment after nerve transection. Acta Neuropathologica. 1997 Mar 19;93(3):252–9.

25. Nocera G, Jacob C. Mechanisms of Schwann cell plasticity involved in peripheral nerve repair after injury. Cell Mol Life Sci [Internet]. 2020 Apr 10 [cited 2020 Jun 29]; Available from: http://link.springer.com/10.1007/s00018-020-03516-9

26. Henzi A, Senatore A, Lakkaraju AK, Scheckel C, Mühle J, Reimann R, et al. Dimeric prion protein ligand activates Adgrg6 but does not rescue myelinopathy of PrP-deficient mice. bioRxiv. 2020 Jan 1;2020.07.07.191452.

27. Berg A, Zelano J, Pekna M, Wilhelmsson U, Pekny M, Cullheim S. Axonal Regeneration after Sciatic Nerve Lesion Is Delayed but Complete in GFAP- and Vimentin-Deficient Mice. Phillips W, editor. PLoS ONE. 2013 Nov 1;8(11):e79395.

28. Triolo D. Loss of glial fibrillary acidic protein (GFAP) impairs Schwann cell proliferation and delays nerve regeneration after damage. Journal of Cell Science. 2006 Oct 1;119(19):3981–93.

29. Hagemann TL, Gaeta SA, Smith MA, Johnson DA, Johnson JA, Messing A. Gene expression analysis in mice with elevated glial fibrillary acidic protein and Rosenthal fibers reveals a stress response followed by glial activation and neuronal dysfunction. Human Molecular Genetics. 2005 Aug 15;14(16):2443–58.

30. Messing A, Head MW, Galles K, Galbreath EJ, Goldman JE, Brenner M. Fatal encephalopathy with astrocyte inclusions in GFAP transgenic mice. Am J Pathol. 1998 Feb;152(2):391–8.

31. Messing A, Brenner M, Feany MB, Nedergaard M, Goldman JE. Alexander Disease. Journal of Neuroscience. 2012 Apr 11;32(15):5017–23.

32. Chinthaparthi S, McDonald K, Davis L. A case of Alexander disease with peripheral neuropathy associated with a novel mutation (P3.326). Neurology. 2018 Apr 10;90(15 Supplement):P3.326.

