## Supplementary figures and images for "The prion protein is not required for peripheral nerve repair after crush injury"

### Supplementary Figure 1

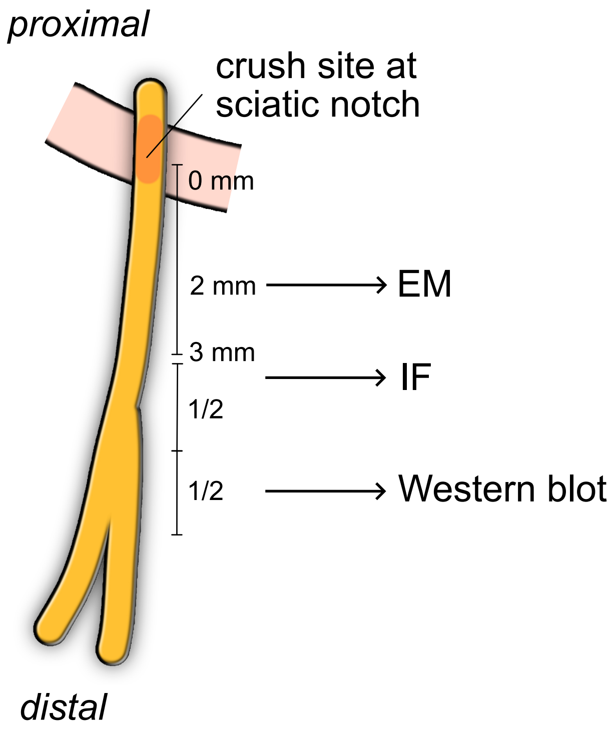
