## Supplementary Figure 2 for "The prion protein is not required for peripheral nerve repair after crush injury"

### Figure 1c

5 d.p.c. c-Jun

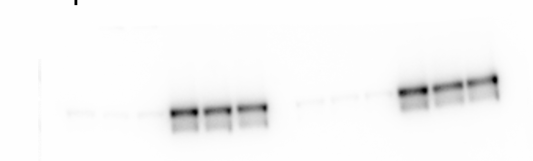

5 d.p.c. Calnexin

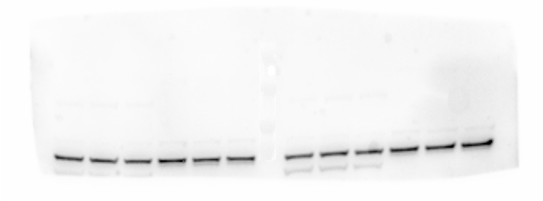

10 d.p.c. c-Jun

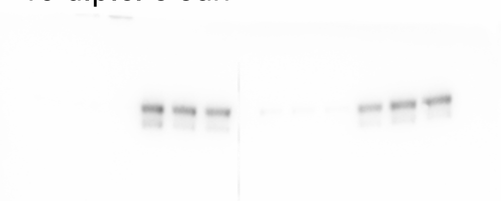

10 d.p.c. Calnexin

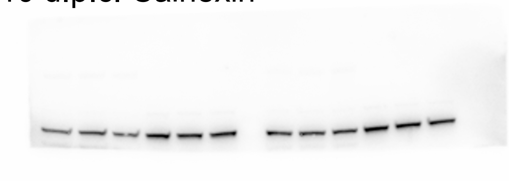

### Figure 1e

blotting with GFAP antibody was performed after c-Jun without stripping

16 d.p.c.GFAP

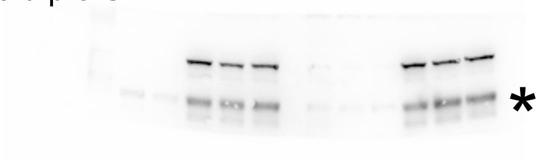

16 d.p.c. Calnexin

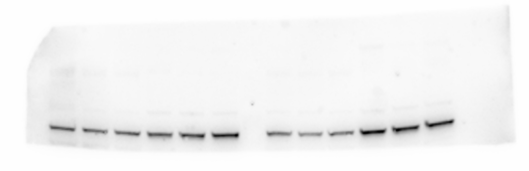

### Figure 1g

GFAP uninjured nerves

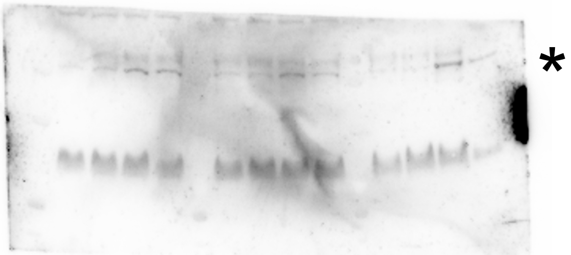

Calnexin uninjured nerves

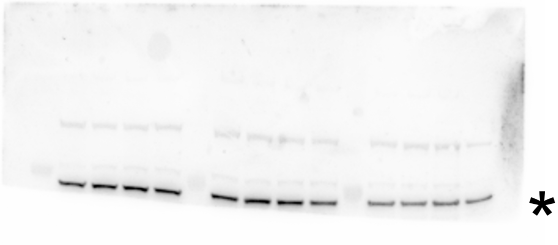

POM2 blotting after GFAP without stripping

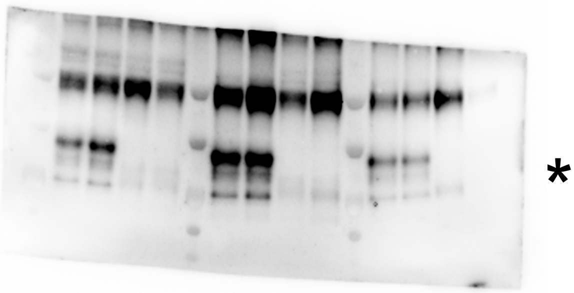

### Figure 1a

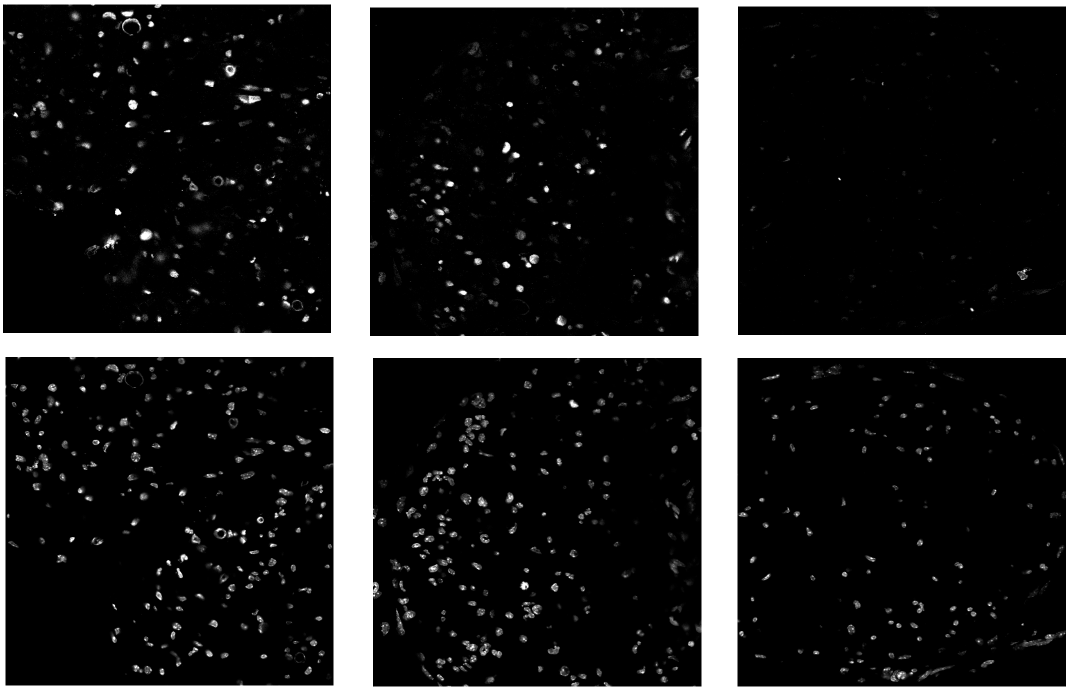

### Figure 2a

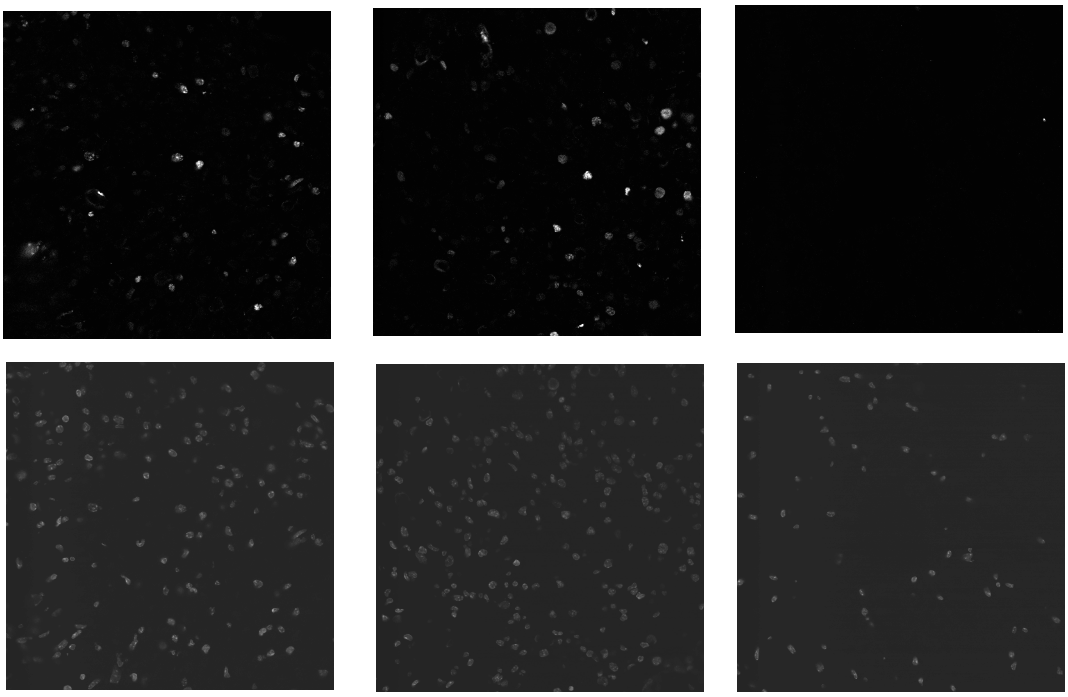

### Figure 1g

Images pseudocolored and channels merges with ImageJ. Correspond to original microscopic images, no adjustment of brightness or contrast

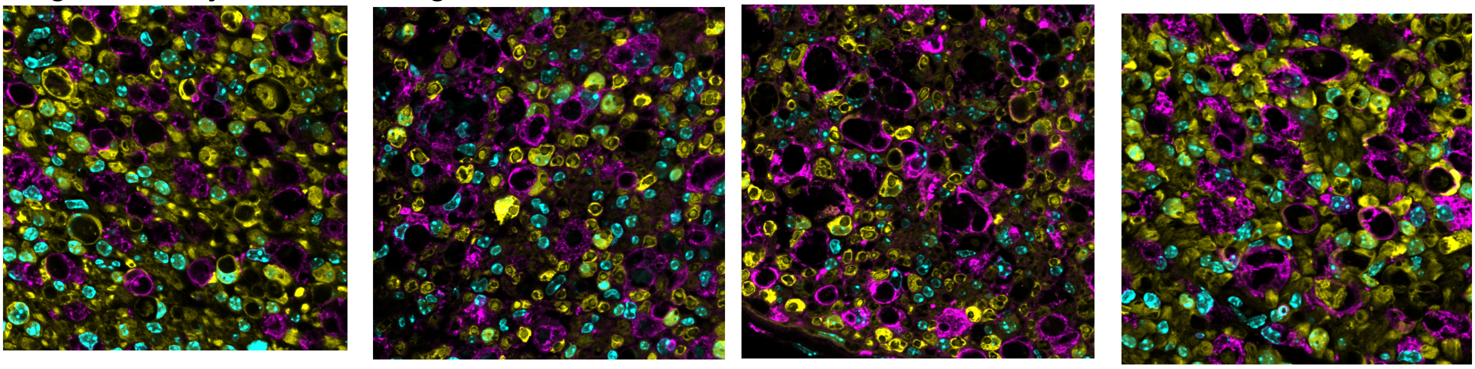
